# Have protein-ligand cofolding methods moved beyond memorisation?

**DOI:** 10.1101/2025.02.03.636309

**Authors:** Peter Škrinjar, Jérôme Eberhardt, Gerardo Tauriello, Torsten Schwede, Janani Durairaj

## Abstract

Deep learning has driven major breakthroughs in protein structure prediction, however the next critical advance is accurately predicting how proteins interact with small molecule ligands, to enable real-world applications such as drug discovery. Recent cofolding methods aim to address this challenge, but evaluating their performance has been inconclusive due to the lack of relevant bench-marking datasets. Here we present a comprehensive evaluation of four leading all-atom cofolding methods using our newly introduced benchmark dataset Runs N’ Poses, which comprises 2,600 high-resolution protein-ligand systems released after the training cutoff used by these methods. We demonstrate that current cofolding approaches largely memorise ligand poses from their training data, hindering their use for *de novo* drug design. With this assessment and benchmark dataset, we aim to accelerate progress in the field by allowing for a more realistic assessment of the current state-of-the-art deep learning methods for predicting protein-ligand interactions.

## 1 Introduction

Being able to accurately predict and characterise protein-ligand interactions (PLI) is a cornerstone of modern drug discovery, enabling the identification of small molecules that can modulate protein function with high specificity. The process of discovering and optimising such modulators into viable lead compounds is a critical aspect of early-stage pharmaceutical development and lead optimisation. Structure-based drug design, a component of computer-aided drug design, relies on three-dimensional structures of the target protein and the protein-ligand interface to inform medicinal chemistry decisions [1–3]. Therefore, to achieve accurate predictions of how ligands bind and their binding affinity, a deep understanding of the fundamental principles governing PLIs is essential [4]. A combination of different computational techniques has become indispensable in both industrial and academic drug discovery efforts, helping to reduce costs and accelerate development timelines [5]. However, despite significant advances in computational methods, accurately modeling protein-ligand binding at atomic resolution remains a significant challenge [6, 7].

Over the years, there have been numerous community initiatives to assess the state of the field in predicting the structure of ligand poses binding to proteins. Recently, CASP15 had a pilot experiment in the evaluation of protein-ligand complex prediction [8], inspired by previous efforts such as Teach Discover Treat (TDT) [9], Continuous Evaluation of Ligand Prediction Performance (CELPP) [10], Drug Discovery Data Resource (D3R) [11–14], and Community Structure–Activity Resource (CSAR) [15, 16], with the added complexity that the prediction task consisted of predicting both the structure of the receptor protein as well as the position and conformation of one or more ligands. Conclusions from CASP15 [17] and parallel benchmarking efforts [18] were threefold: (1) The joint protein-ligand prediction task requires accuracy metrics that take both protein and ligand positioning into account, such as binding-site superposed RMSD and local difference distance test (LDDT) of protein-ligand interactions (LDDT-PLI) [17]; (2) Experimental quality is a more acute issue for PLI prediction compared to protein structure prediction, due to the atom-level reliability required and the fact that many PDB entries fail to meet even relaxed X-ray quality standards [18, 19] to enable their use as reliable ground truth; and (3) Many complexes with ligands released in the PDB are either very similar to previously released complexes or not relevant for drug discovery applications, or both. While CASP16 addressed part of the latter issue by soliciting unreleased PLI complexes from pharmaceutical companies, the scarcity of targets and the availability of very similar templates in the PDB make it difficult to draw conclusions specifically about deep learning methods that qualify for this task, most of which also did not participate.

Deep learning methods such as AlphaFold2 [20] have revolutionised protein structure prediction, inspiring the development of all-atom cofolding methods like RosettaFold All-Atom [21], AlphaFold3 [22], Chai-1 [23], Boltz-1 [24], Protenix [25], and NeuralPLexer3 [26]. While these methods show promise, evaluating their performance is difficult due to the lack of established benchmarking techniques for joint protein-ligand binding in the deep learning domain. Existing benchmarks often fail to account for similarities between the training data and test sets, leading to inflated performance estimates [18, 19, 27, 28]. This is particularly problematic for PLI systems, where the scarcity and bias in experimental structures can exacerbate both the risk of memorisation and our inability to detect it.

To address this, we present a comprehensive benchmark of four leading allatom cofolding deep learning methods with highly similar architectures and training paradigms (AlphaFold3 [22], Chai-1 [23], Protenix [25], and Boltz-1 [24]) on 2,600 high-resolution PLI systems released after their training cutoff (30 September 2021). We explore the impact of training data similarity on prediction accuracy, revealing a critical limitation: current cofolding methods struggle to generalise beyond ligand poses seen in their training data. We also compared the top-ranked and best-scored models, observing a slight performance improvement, yet the overall trend of limited generalisation remained unchanged.

This overfitting issue, independent of model confidence scores, highlights the need for improved architectures and data augmentation strategies. Notably, we observe better generalisation for cofactor, amino acid, and nucleotide analogs, suggesting the importance of abundant training data. We provide a benchmark dataset (Runs N’ Poses) with varying degrees of training set similarity, along with resources to analyse the relationship between data leakage and performance. With this work, we aim to facilitate progress and enable identification of genuine breakthroughs in the critical field of protein-ligand complex prediction.

## 2 Results

### 2.1 Limited generalisation of cofolding methods across protein-ligand complex space

We aimed to explore the dependence of prediction accuracy on the similarity of the target complex to the training set for the four deep learning cofolding methods, based on our benchmark dataset of 2,600 systems (covering 3,047 non-ion non-artifact ligands). Our analysis shows a clear correlation between the accuracy of the top-ranked predicted model for each method and the similarity of the reference system to the training set, depicted in Figure 1A.

**Fig. 1.**
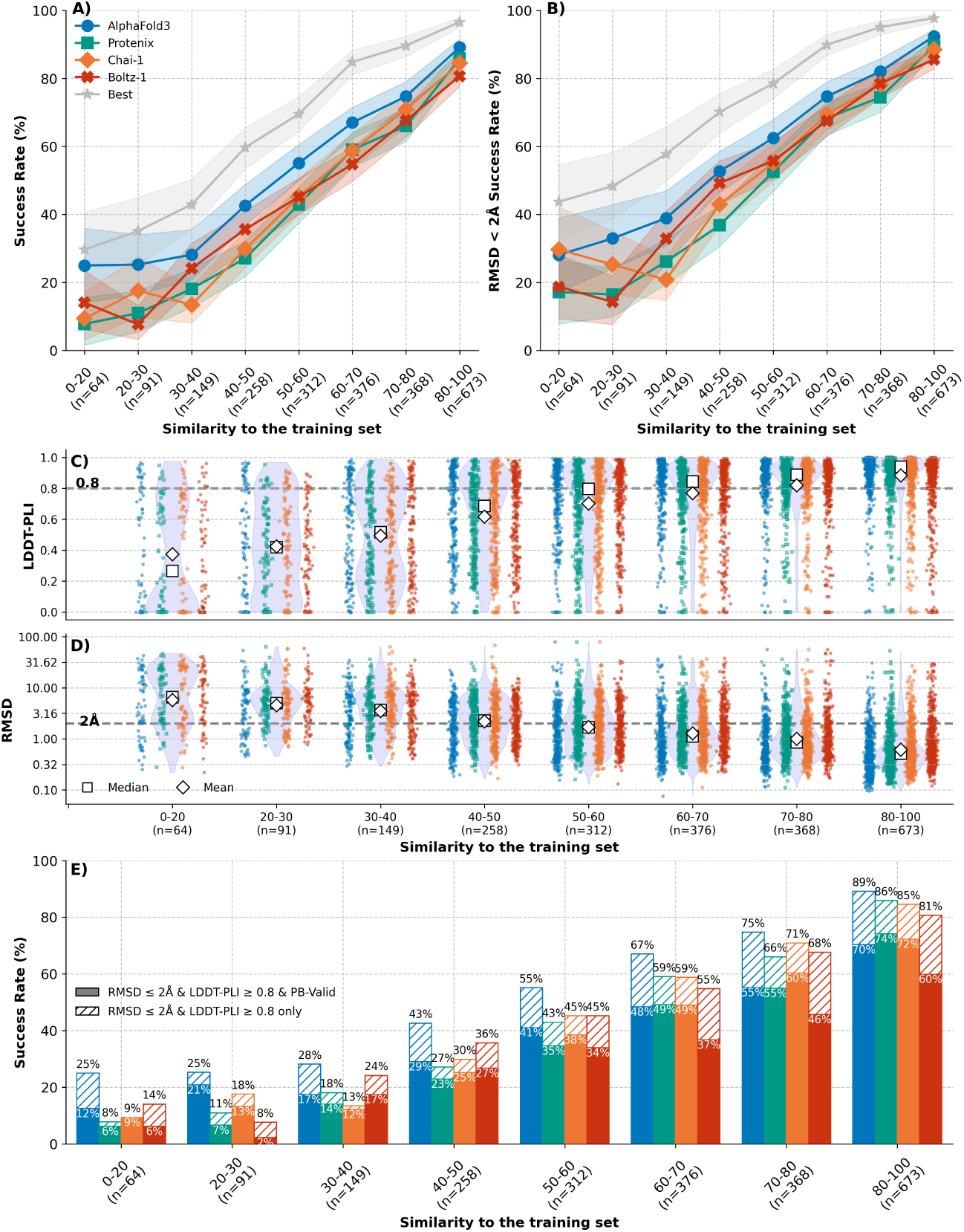
Prediction accuracy vs. training set similarity. **A)** The success rate, defined as the percentage of system ligands with *<*2Å RMSD and *>*0.8 LDDT-PLI, for all common subset system ligands in each similarity bin across the four methods and when selecting the best-scored model from the four top-ranked. Shaded regions correspond to the 95% confidence interval, calculated from 1,000 bootstrap samples for each bin and method. **B)** same as A but with success rate defined only by <2Å RMSD. **C), D)** The distribution of LDDT-PLI and RMSD values respectively for all methods displayed as a violin plot, with each individual method being shown as coloured scatter points. **E)** Same as panel A with added physical validity checks. The striped bars show the share of predictions of each method that have *<*2Å RMSD and *>*0.8 LDDT-PLI and the solid bars show the subset that in addition have valid geometries and energies, i.e., pass all PoseBusters tests and are therefore ‘PB-Valid’.

Here, we define the similarity to the training set as the product of binding pocket coverage [28] and Combined Overlap Score (SuCOS) of the ligand pose, which measures the extent to which ligand poses overlap in both volume and placement of chemical features [29, 30]. Thus, a training system is only considered similar if it has a similar ligand pose in a similar protein pocket to the test system (see Section 2.4 for further details on the protein and ligand similarity metrics).

Moreover, in classical docking benchmark studies, success is typically defined solely by ligand RMSD (*<* 2Å), as the protein structure is generally treated as rigid and assumed to be correct. However, for cofolding methods, it is essential to assess the accuracy of both the ligand and the protein structure. Therefore, we define the success rate as a combination of LDDT-PLI (*>* 0.8) [17] and ligand pose RMSD (*<* 2 Å) (see Section 5.3 for further details on this choice). Figure 1B, where success is defined solely on RMSD (*<* 2Å), confirms that our success rate criteria do not influence the observed trend of limited generalisation. As shown in Figure 1C and D, we observed that LDDT-PLI and RMSD distributions for each method consistently correlate with the similarity to the training set (Figure 1C, 1D).

Finally, Figure 1E shows the fraction of predictions passing physical validity checks as defined by the PoseBusters (PB) suite [19]. The most problematic is the minimum distance to protein check, which passes if the distance between protein-ligand atom pairs is larger than 0.75 times the sum of the pairs van der Waals radii, with 22.6% of predicted models failing this check across the four methods. Failing this check suggests physically implausible positioning of atoms that would lead to steric clashes between the protein and the ligand. The remaining checks are passed by *>*95% of predicted models.

### 2.2 Model and data choices lead to minimal performance improvement

In order to directly compare methods to each other, Figure 1 focused on the common subset of systems that were predicted by all four methods using their default settings, ignoring cases where certain methods fail to produce an output due to varying steps in their data processing and prediction pipelines (see Table 1 and Section 5.2 for statistics on the method-specific failures). Although all four methods exhibit comparable correlations of accuracy with their training set similarity, AlphaFold3 demonstrates a slight performance advantage over the others, despite all methods sharing similar architectures and training paradigms. This advantage may stem from differences in training times, dataset sampling strategies, validation sets, and other methodological choices between the four methods. However, the methods seem to have some level of complementarity, as indicated by the gray line representing the best-scored of the top-ranked models across all four methods (Figure 1A and 1B).

**Table 1.** Counts of the full Runs N’ Poses dataset and the obtained predictions from the methods benchmarked. AF3 = AlphaFold3, AF3-NT = AlphaFold3 run without templates, RFAA = RosettaFold-All-Atom.

| Method | PDB IDs | Systems | All Ligands | Proper ligands | Multi-ligand systems | Multi-protein systems |
| --- | --- | --- | --- | --- | --- | --- |
| Post 30 September 2021 |  |  |  |  |  |  |
| Total | 2,585 | 2,600 | 4,282 | 3,047 | 401 | 790 |
| AF3 | 2,396 | 2,409 | 3,860 | 2,710 | 296 | 738 |
| Protenix | 2,310 | 2,323 | 3,726 | 2,609 | 281 | 702 |
| Chai-1 | 2,280 | 2,292 | 3,683 | 2,572 | 275 | 685 |
| Boltz-1 | 2,126 | 2,139 | 3,372 | 2,400 | 256 | 626 |
| AF3-NT | 2,398 | 2,411 | 3,861 | 2,711 | 295 | 738 |
| Boltz-1x | 2,078 | 2,091 | 3,322 | 2,338 | 242 | 608 |
| RFAA | 1,759 | 1,769 | 2,718 | 1,946 | 173 | 507 |
| Post 1 June 2023 |  |  |  |  |  |  |
| Total | 1,025 | 1,028 | 1,685 | 1,187 | 142 | 415 |
| Boltz-1 | 841 | 844 | 1,322 | 933 | 87 | 347 |
| Boltz-2 | 868 | 871 | 1,396 | 961 | 88 | 361 |

Supplementary Figure S1 shows the same panels calculated on all predictions, and in addition shows the results from (1) AlphaFold3 run without templates (light blue), (2) Boltz-1x (pink), and (3) RoseTTAFold-All-Atom [21] (purple). AlphaFold3 with and without templates does not show any significant differences; this is expected since ligand information from the template is not utilised by the method. Boltz1x does not show differences in success rate compared to Boltz-1 v0.4 but has a boost in PoseBusters physical validity checks, with all predicted systems passing the minimum_distance_to_protein check, and outperforming the other methods as previously reported [24]. RoseTTAFold-All-Atom has a different architecture and was run with only 1 model output instead of 25 and thus is not directly comparable, however this default setting performed poorly compared to the others.

Since the creation of Runs N’ Poses, Boltz-2 [31] has been released which was trained with a cutoff of 2023-06-01, thus using a full two years more PDB data than the other methods. While this makes the new method not directly comparable to the others presented in this manuscript, it can still be assessed using the similarity to the training set. We compared Boltz-2 performance on Runs N’ Poses systems released after the 2023 training cutoff, as a function of the similarity to the PDB up to this point, to the Boltz-1 performance with similarities calculated according to the Boltz-1 training cutoff. As seen in Figure 2A, the additional data used for training (over 25k PDB entries) increases the number of cases in the higher similarity bins but does not seem to help with generalisation to unseen data.

**Fig. 2.**
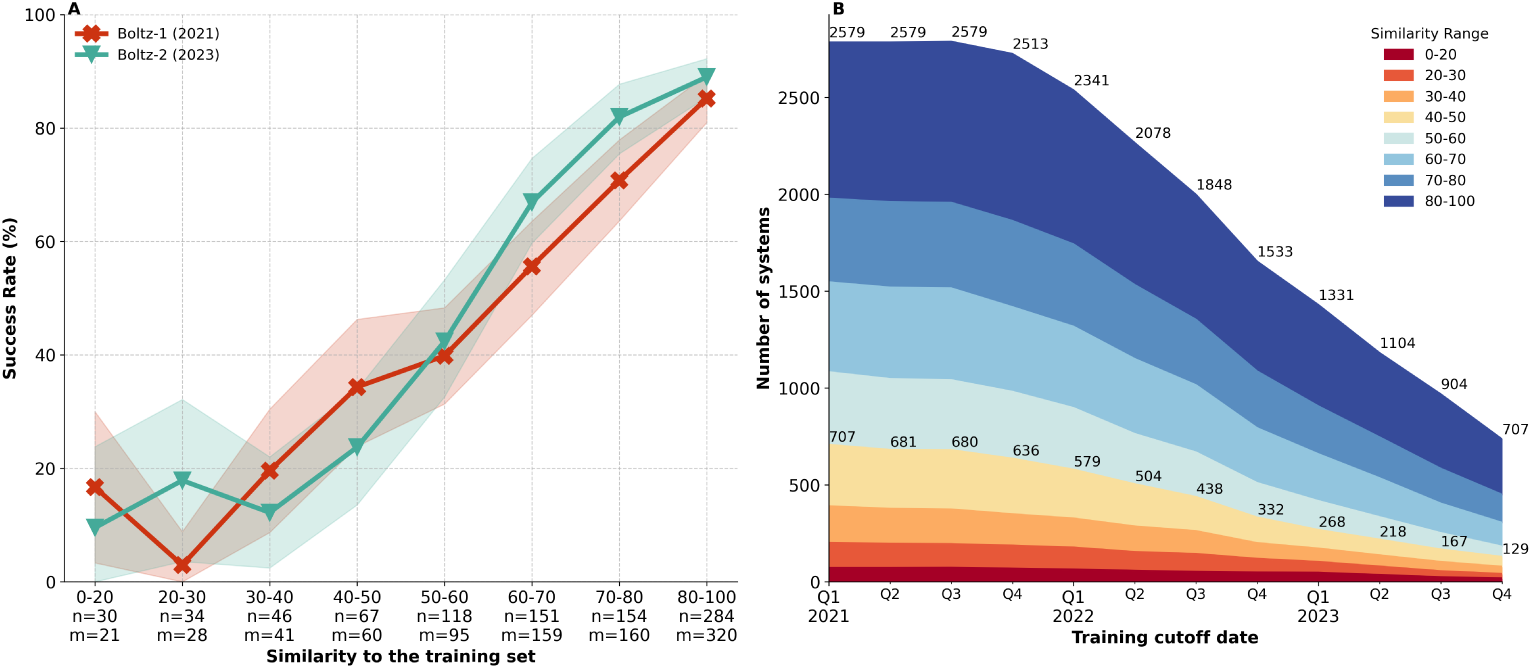
Benchmarking with different training cutoffs. **A)** The success rate, defined as the percentage of system ligands with *<*2Å RMSD and *>*0.8 LDDT-PLI, for systems released after the Boltz-2 training cutoff when predicted by Boltz-1 (green) and Boltz-2 (red) for each similarity bin. The similarity bins are calculated using the 30 September 2021 training cutoff for Boltz-1 (counts labelled as *n*), and using the 1 June 2023 training cutoff for Boltz-2 (counts labelled as *m*). **B)** Training set similarity distributions for different training cutoffs. The counts of systems with *<*50 similarity and the total number of systems released after the cutoff are labelled.

Figure 2B demonstrates the amount of systems available in Runs N’ Poses for various training cutoffs as well as their distribution across the different similarity bins. While there is still a reasonable number of cases for benchmarking even post 2023, there is a significant decrease of cases in the more difficult bins, lowering the capability to perform granular and confident analysis of memorisation and generalisability.

### 2.3 Impact of ligand and complex prevalence

Further stratification reveals slightly improved performance on ligands with significant amounts of training data (Figure 3A). This includes cofactors and their analogs, as these are found across a wide variety of protein families and thus also across protein-ligand structures and pockets in the PDB. It also includes analogs of amino acids and nucleotides, likely a benefit of training on protein-protein complexes and protein-nucleotide complexes. To set these apart from more drug-like molecules, we classify them based on their frequency of occurrence in the PDB, independent of the protein and pocket. Specifically, for each ligand in the test set we counted the number of times any analogous ligand appears in any system in the training data. We define analogous ligands as those with *>*0.9 ligand RDKit topological fingerprint Tanimoto similarity [32] to the query ligand. Supplementary Figure S2 depicts molecules with more than 100 analogous occurrences, hereafter referred to as *prevalent*. Removing these ligands from consideration, i.e keeping only *distinct* ligands, shows a more linear trend of similarity to success rate (Figure 3B), demonstrating that the successes in the lower similarity bins are primarily due to such cases. This analysis indicates that data scarcity is one of the drivers of memorisation, as drug-like molecules typically do not have a lot of experimental structures resolved in different protein pockets.

**Fig. 3.**
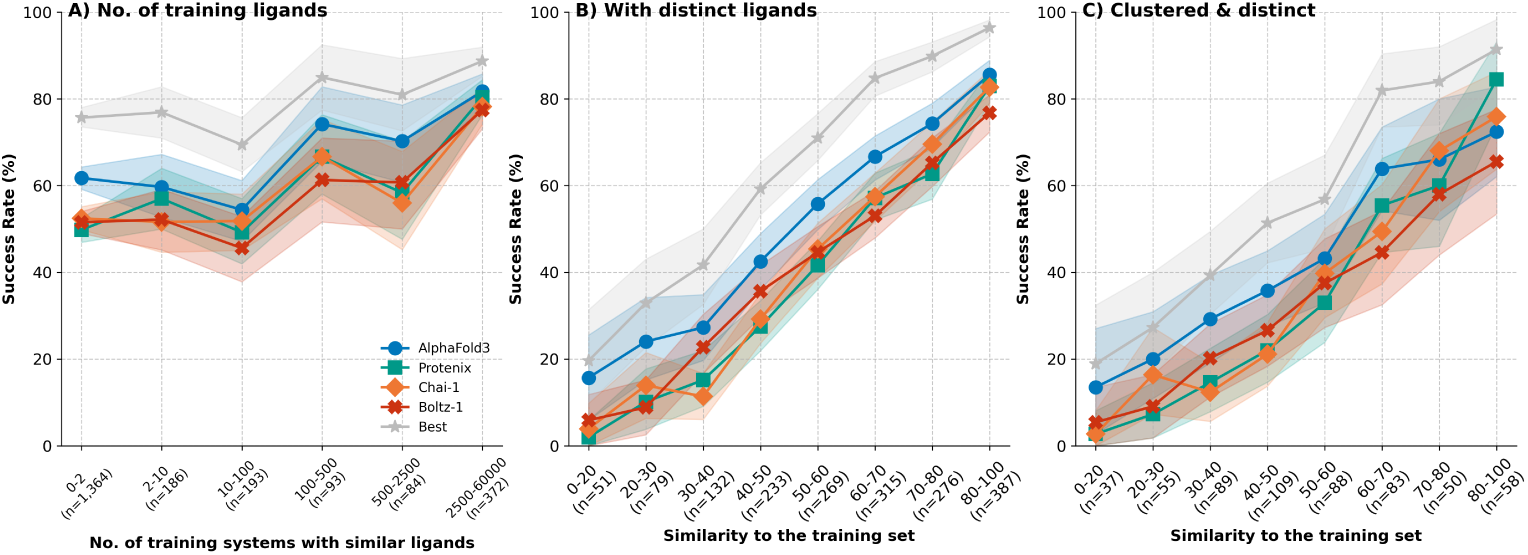
Prediction accuracy for prevalent vs. distinct ligands. The success rate, defined as the percentage of system ligands with *<*2Å RMSD and *>*0.8 LDDT-PLI across the four methods and when selecting the best-scored model from the four top-ranked, for **A)** system ligands with differing numbers of analogous ligand training systems., **B)** only system ligands having *<*100 analogous ligand training systems across each similarity bin, and **C)** only cluster representatives from B across each similarity bin. Shaded regions correspond to the 95% confidence interval, calculated from 1,000 bootstrap samples for each bin and method.

We verified that the observed trend is not driven by an over-representation of certain protein families, which could bias the results. Figure 3C displays the same trend correlating success vs. training set similarity only for the cluster representatives of distinct ligand systems, based on graph community clustering at SuCOS-pocket similarity *>* 50. This leads to a total of 789 clusters in the common subset out of 1,017 total clusters. Supplementary Table S1 details the sizes, cluster representative details, and similarity bin coverage of the ten biggest clusters. The first cluster consists of 171 crystal structures of SARS-CoV2 main protease (MPro) in complex with different small molecules from independent efforts such as the SARS-CoV2 moonshot series [33] and others [34], while the second consists of 136 protein kinases complexes with various inhibitors. Thus SuCOS-pocket clustering puts together protein-ligand complexes where similar molecules bind similar proteins in the same pocket and with similar three-dimensional poses. For example, the HCov-NL63 MPro complexes in cluster #4 have different binding modes than those in cluster #1. While Figure 3C shows that the memorisation trend is independent of redundancy, such congeneric series, i.e. ligands sharing the same core structure and differing only in some substituent groups, are of high relevance to drug discovery applications and accuracy across these series is an important target to aim for.

We also show that the methods are not biased in performance towards smaller or larger ligands, as seen in Supplementary Figure S3A and S3B. Although we observed a slight correlation between success rate and the binding pockets size (Supplementary Figure S3C), the lack of generalisation trend is still seen across different binding pocket sizes (Supplementary Figure S3D-H). Moreover, looking at different subsets of the dataset, such as single-ligand, multi-ligand, and multi-protein systems only, shows again the same pattern, demonstrating the effectiveness of our dataset for benchmarking these sub-tasks (Supplementary Figure S3I-K).

### 2.4 Ligand and protein similarity metrics for detecting data leakage

To explore how much the memorisation trend is influenced by similarity at protein, pocket and ligand levels, Figure 4A and 4B shows how protein sequence identity, pocket coverage, ligand SuCOS, and ligand fingerprint similarity (using the RDKit topological fingerprint and Morgan fingerprint with radius 1 and 2048 bits), evolve across similarity bins. This reflects the relationship between each system and its closest training system as determined by SuCOS-pocket similarity. Here, the relationship between the SuCOS score and pocket coverage becomes clearer, as the lowest bin (SuCOS-pocket similarity between 0 and 20) includes examples with high SuCOS but low pocket coverage, and vice versa. In contrast, the protein sequence identity and the ligand fingerprint similarity vary more broadly, but also tend to increase with increasing SuCOS-pocket similarity. We conclude that previous similarity metrics and thresholds used for validating cofolding methods’ performance are insufficient to detect data leakage for the PLI prediction task. The commonly used 40% sequence identity threshold misses numerous cases of proteins from the same family sharing the same fold, pocket and binding mode, and 85% Morgan fingerprint similarity threshold (used for validation of AlphaFold3 [22]) misses cases of ligands binding with highly similar poses to highly similar proteins and pockets, as exemplified in Figure 4C and 4D respectively.

**Fig. 4.**
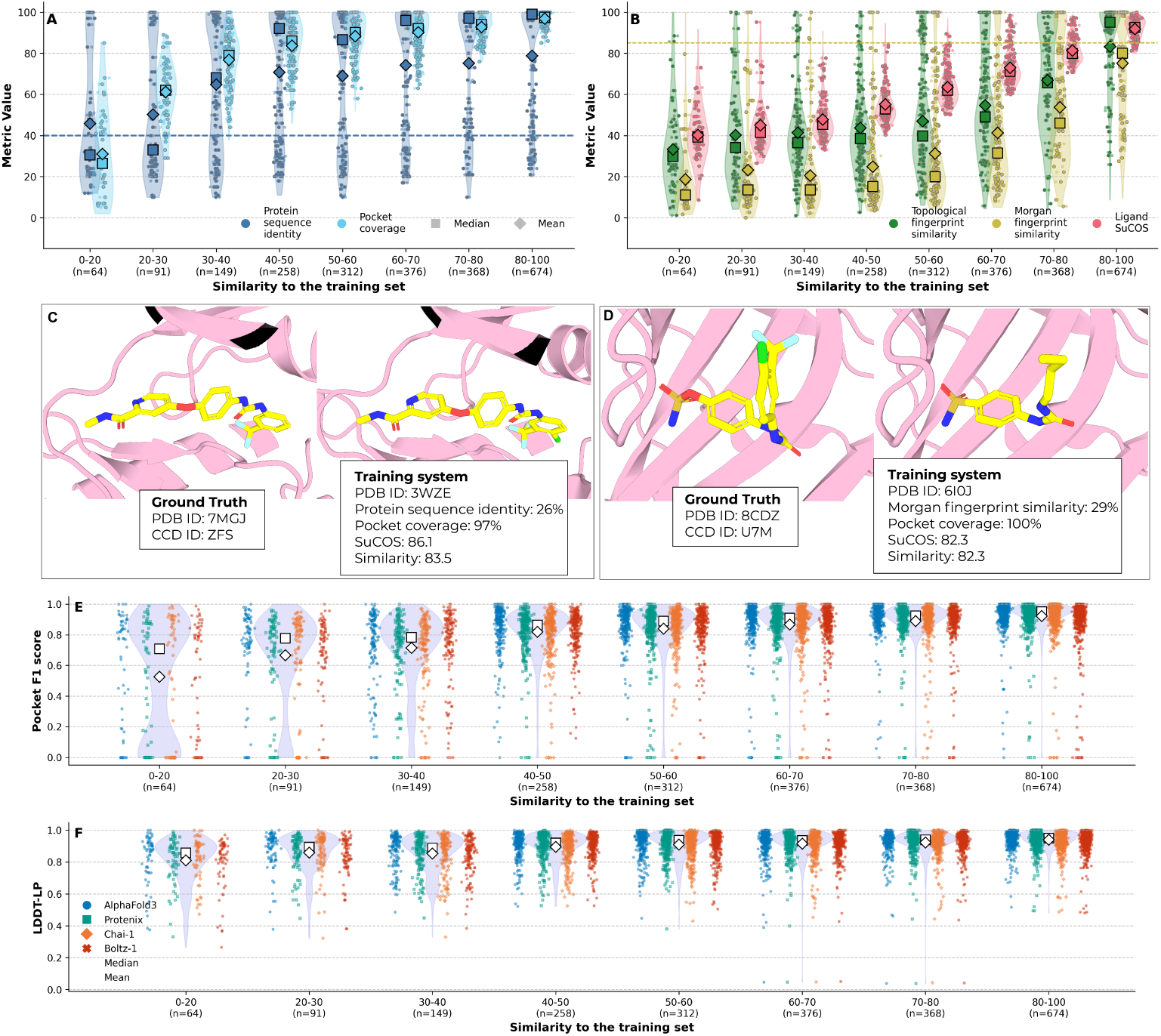
Comparison with other similarity and accuracy metrics. **A)** The distribution of protein sequence identity and pocket coverage (both shown as percentage) for the closest training system found by SuCOS-pocket similarity. The dashed line indicates the protein sequence identity threshold used in validation of AlphaFold3. **B)** The distribution of ligand topological fingerprint Tanimoto similarity, ligand Morgan fingerprint Tanimoto similarity (radius=2, nbits=2048), and ligand SuCOS similarity for the closest training system found by SuCOS-pocket similarity. The dashed line marks the ligand Morgan fingerprint Tanimoto similarity threshold used in validation of AlphaFold3. **C)** An example complex with low sequence identity and high SuCOS-pocket similarity to the closest training system. **D)** An example complex with low Morgan fingerprint similarity and high SuCOS-pocket similarity to the closest training system. **E)** F1-score distribution for recovering binding pocket residues across similarity bins for all methods (gray violin) and each method separately (coloured scatter). **D)** Distribution of LDDT-LP values across similarity bins for all methods (gray violin) and each method separately (coloured scatter). Percentage of systems with LDDT-LP *>* 0.8, for all systems: AlphaFold3: 92%, Protenix: 91%, Chai-1: 90%, Boltz-1: 91% For cluster representatives (gray outlines): AlphaFold3: 87%, Protenix: 83%, Chai-1: 82%, Boltz-1: 83%

Interestingly, already from the third bin (SuCOS-pocket similarity between 30 and 40) we see that the protein and pocket of a test system share more than 60% sequence identity and 80% pocket coverage to the closest training system on average. This is further supported by the Figure 4E, which shows the F1-score distribution for recovering binding pocket residues across different similarity bins. While there is a correlation with the similarity to the training set, finding the correct pocket does not seem to be the limiting factor for the inaccurate complex predictions. Thus, the trend that we observe is not a case of lack of out-of-distribution (OOD) generalisation for lesser-studied proteins or novel pockets and more likely arises from other factors, such as the difficulty in generalising to unseen ligand poses within otherwise familiar pockets.

To illustrate binding pocket conformation prediction, Figure 4F shows the persimilarity bin distribution of LDDT-LP, i.e. the accuracy of the relative positioning of residues in the binding pocket. Although there is some increasing trend across the bins, the majority of predictions have an LDDT-LP *>* 0.8, indicating that the protein is well-modelled. However, these results do not guarantee that the predicted protein models could be used for downstream docking applications, as previous studies showed that even LDDT-LP values higher than 0.9 do not detect atom-level conformational changes which lead to unsuccessful rigid docking [18].

Figure 5 shows some examples of successful and failed predictions from different similarity bins along with the ground truth complex, and the closest training system found based on the SuCOS-pocket similarity score. As the similarity to the training systems increases, we observe, as expected, a growing overlap in terms of molecular shape and common substituents between the target ligand and the one found in the closest training system. However, in some cases, important chemical features are overlooked during inference, as exemplified in Figure 5E. Despite being explicitly defined in the input SMILES, all the methods place the 5-chlorobenzofuran-3-aminomethyl group with incorrect chirality, perhaps mimicking the 5-chloro-2-methoxyphenyl group found in the closest training system. Regarding proteins, globally, we observe that their structures and binding pockets are accurately predicted across all similarity bins, as depicted in 4D. This is not always the case, as Figure 5C shows an example where also the protein is mispredicted - the ground truth is a protein kinase in inactive DFG-out conformation [35], however the predicted structure adopts a DFG-in conformation. Although Figure 5 provides some useful visual cues on when these current methods are likely to perform well, it also paints a clear picture of their inherent limitations in generating accurate predictions for drug-like molecules beyond the training data.

**Fig. 5.**
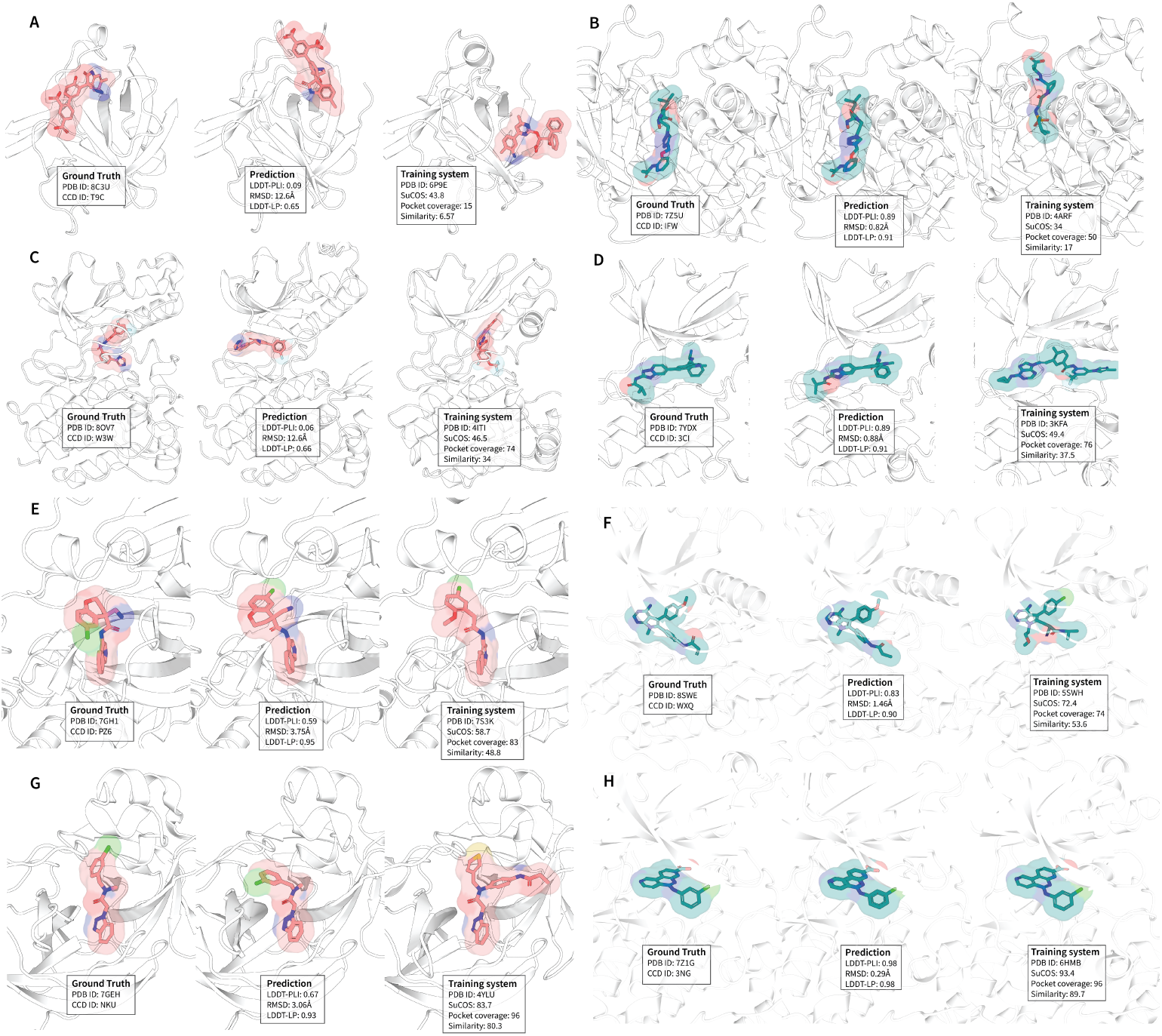
Examples of incorrect and correct predictions across different similarity bins. Predicted models from AlphaFold3 (middle of each panel) compared to the ground truth experimental structure (left) and the closest training system by SuCOS-pocket similarity (right). For each panel, the PDB ID and CCD code of the ground truth, the RMSD, LDDT-PLI and LDDT-LP of the prediction, and the PDB ID, SuCOS score, pocket coverage (percentage), and SuCOS-Pocket similarity score of the closest training system are labelled. We depict an incorrect prediction (red, left panels) and a correct prediction (green, right panels) from similarity bins 0-20 (**A-B**), 30-40 (**C-D**), 40-60 (**E-F**) and 80-90 (**F-G**).

### 2.5 Room for improvement in sampling and ranking of ligand poses

To de-correlate the effect of predicting poses versus ranking them, we look at the accuracies for all 25 models (5 seeds, 5 diffusion samples) for each system across the four methods. Figure 6A-D depicts, for each method and SuCOS-pocket similarity bin, the success rates of the best-scored (i.e closest to the ground truth), worst-scored (i.e furthest from the ground truth), a randomly picked model from the 25, and the top-ranked model selected based on each method’s ranking scores. We also look at the distribution of top-ranked, best-scored and randomly picked models across the 5 seeds, to demonstrate the impact of using multiple seeds. While the top-ranked model is better than the worst-scored model generated (black vs. red), there is still a gap to the best possible model (in blue) and it is usually not much better than random selection (in green). Additionally, we observe that using multiple seeds is currently not useful at inference time for any of the methods. Although more accurate ligand poses are sampled with 5 seeds (blue vs. yellow), the ranking picks similar models when using 1 or 5 seeds (as seen by the black line overlapping with the gray and purple lines). Nevertheless, sampling the correct pose continues to be the main bottleneck in improving the accuracy, since even the best-scored poses do not flatten the curve.

**Fig. 6.**
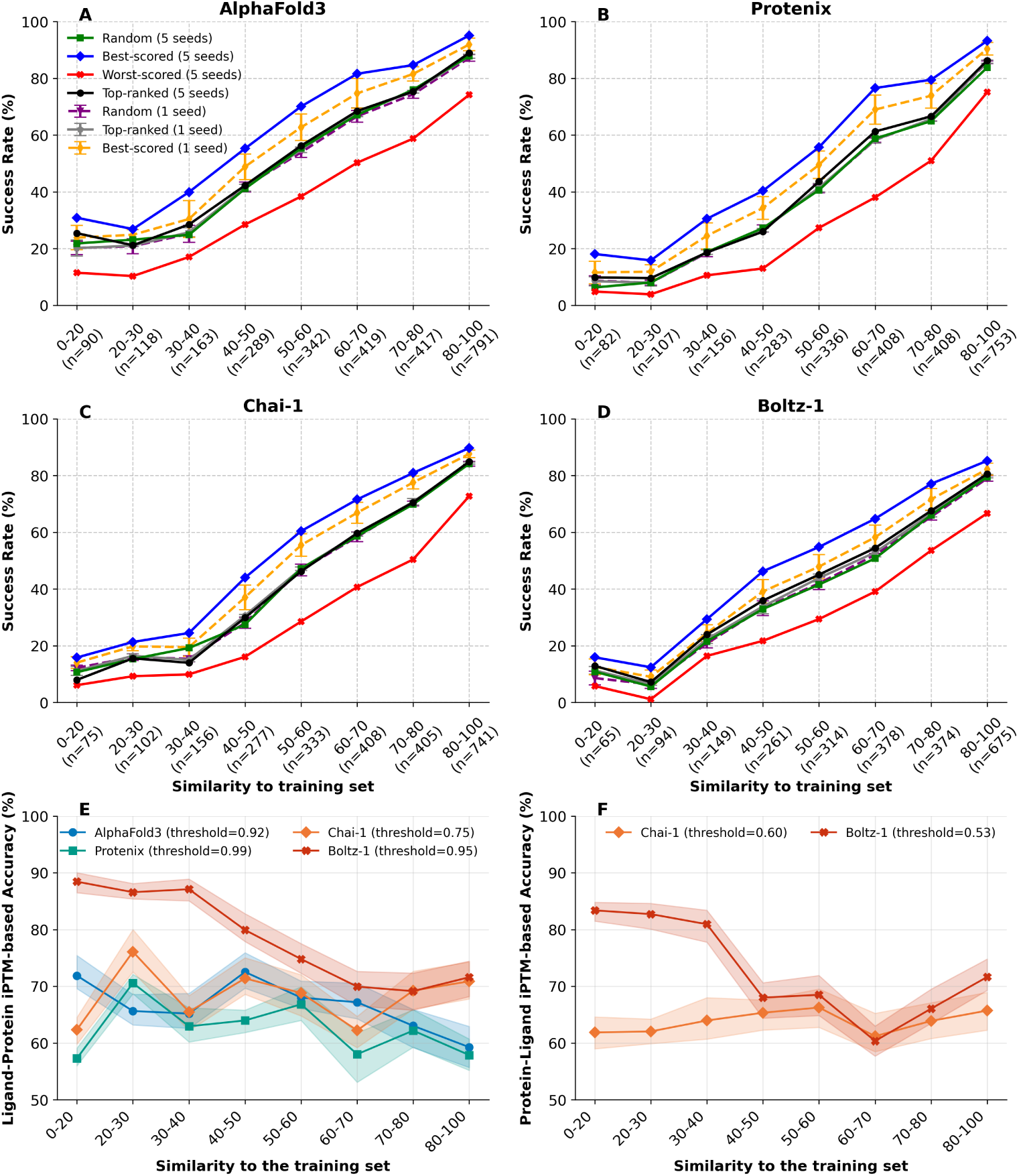
Confidence metrics within and across systems. **A-D)** For each method, the success rate across similarity bins for a randomly selected model across 25 (green squares), the best-scored model (blue diamonds, best LDDT-PLI across 25), worst-scored model (red crosses, worst LDDT-PLI across 25), top-ranked model (black circles, with the highest ranking score), top-ranked across 1 seed (gray pentagons, with standard deviation as whiskers across the 5 seeds), randomly selected from 1 seed (purple triangles), and best-scored across 1 seed (yellow diamonds). **E)** Ligand-protein chain-pair iPTM-based success classification accuracy for the four methods. 250 positive and 250 negative systems are sampled for each similarity bin and each method, with 1000 bootstrap estimates shown as shaded areas. **F)** Same as E but using protein-ligand chain-pair iPTM and only for Chai-1 and Boltz-1.

We also investigated the usability of the ligand-protein chain pair iPTM score for differentiating accurate and inaccurate poses across predicted models of different systems. We calculate this score as the average of chain pair iPTM scores across all protein chains to the considered ligand chain. For each method, we independently determine the best threshold as the optimal operating point of the ROC-AUC curve across all predicted systems, where positives are predicted models with LDDT-PLI *>* 0.8 and RMSD *<* 2Å compared to the reference and negatives are predicted models that don’t satisfy this criteria. Thus, this chosen iPTM threshold has the best trade-off between sensitivity and specificity. The optimal threshold is different for each method, with Protenix being the most optimistic (0.99) about its predictions, Chai-1 being the least (0.75), and AlphaFold3 and Boltz-1 having thresholds of 0.92 and 0.95 respectively. Thus, previously established TM-score thresholds cannot be used for model selection, and the confidence scores are not comparable across the four methods.

Figure 6E shows the accuracy across the similarity bins on classifying predicted models as positives or negatives based on the determined iPTM threshold for each method. Equal numbers of positive and negative cases (250 each) were chosen for every bin, to counteract the imbalance across the similarity bins. Overall, the iPTM-based accuracy does not seem to correlate with the training set similarity. Out of the four assessed methods, Boltz-1 shows a clear advantage over the others in accurately classifying their predictions with 70-90% accuracy across the similarity bins, whereas the other methods stay below 75%. Compared to the other methods, Boltz-1 introduced several architectural and training modifications to their confidence model, clearly reflected in improved iPTM-based ranking power.

An additional peculiarity differentiating the four methods is the asymmetrical nature of the chain-pair iPTM matrix for Boltz-1 and Chai-1, where the protein-ligand chain pair iPTM is a different value than the ligand-protein chain pair iPTM. This is not the case for AlphaFold3 and Protenix, both of which have symmetrical matrices due to averaging across protein and ligand chains. The protein-ligand chain-pair iPTM shows lower ranking power (Figure 6F).

## 3 Discussion

Overall, our benchmark shows that the current generation of deep learning cofolding methods still have a long way to go to accurately model unseen protein-ligand complexes. While all methods achieve high accuracy at modelling the protein part, protein-ligand interaction and ligand pose prediction remain the primary challenge. We demonstrate that the performance of current approaches strongly correlates with the similarity to their training data, regardless of the metric used to define success or the subsets considered. Our analysis suggests that both sampling and ranking contribute to the limitations of these methods. In cases where the prediction was not accurate, the correct pose is either not sampled at all or, when it is, the method fails to select it during ranking, further highlighting the need for improvement in both aspects.

The results on prevalent ligands found in multiple training systems across diverse pockets indicate that data scarcity remains a major challenge for these methods. Given the vastness of chemical space, we are far from the optimal data regime required for deep learning models to generalise effectively. Data augmentation strategies such as cross-docking [36, 37], direct usage of underlying electron densities [38–40], or using small molecule data in isolation [41, 42] may help improve the diversity and coverage of training data across chemical space as well as across different pairs of pockets and ligands. In addition, a large number of experimentally resolved protein-ligand complexes, binding assays useful for cross-docking, and negative binding data are available within pharmaceutical companies and closed-source settings. A collective effort to make such industrial data available for training, perhaps through infrastructures for distributed privacy-preserving federated learning such as the MELLODDY project [43], or large-scale targeted experimental structure determination as planned by the OpenBind consortium [44], could greatly improve the data scarcity situation.

However, even with increased dataset diversity, it remains unclear whether the optimal architecture has been developed for this task or if current methods would fully exploit these advancements. For example, the generalisability of Boltz-2 did not discernibly improve compared to Boltz-1, despite being trained on more data (Figure 2). Drawing parallels with the protein structure prediction task, AlphaFold2 [20] benefited from co-evolutionary signals extracted from billions of sequenced proteins to learn an approximate energy landscape [45, 46] and truly achieved generalisable performance also on difficult proteins without homologous structures in the PDB, with the caveat of having poorer performance on orphan proteins without such co-evolutionary information [47]. All-atom cofolding methods do not currently rely on any such external non-structural data for encoding and representing ligand interactions. Incorporating physics-based terms to more accurately model protein-ligand interactions, potentially from simulations, conformational ensembles, or other sources, are likely needed to achieve more exciting results in this field. Our results are also in line with recent work demonstrating, through adversarial examples, significant divergences from expected physical behaviors in protein-ligand predictions [48]. This highlights the need to integrate robust physical and chemical priors to enhance method generalisation. In future work, we aim to include template-based and physics-based docking baselines, using similar systems from before the training cutoff as templates for homology modelling. While this template reliance may still lead to a marginally increasing trend across similarity bins, we expect this baseline to have a much flatter slope in Figure 1, as physics-based docking is typically not reliant on training sets [18].

All in all, our findings stress the importance of specialised benchmark datasets and evaluation techniques tailored to deep learning PLI prediction. We observed that most released high-quality complexes have a high similarity to the training set (Figure 1, 75% system ligands have *>*50 SuCOS-pocket similarity), demonstrating that difficult test systems consisting of novel molecules or binding pockets, both highly relevant to drug discovery and design, are underrepresented in general, and the commonly-used time split approaches would not capture these. In fact, 87% of the complexes in the commonly used PoseBusters benchmark dataset [19] have a SuCOS-pocket similarity *>*50 to the training set, explaining the high performance scores typically obtained by these methods on this set.

In fact, out of the four benchmarked methods only the authors of Protenix list memorisation artifacts as an expected limitation, further underscoring the difficulty in uncovering such biases without extensive benchmarking combining measures of prediction performance with measures of data leakage. We demonstrate the pitfalls of commonly used similarity thresholds such as 40% sequence identity, proving them to be insufficient to assess method generalisation. While SuCOS similarity proved most relevant for this analysis as it combines both volume and chemical features of the small molecule, it would also be interesting to disentangle these two aspects and inspect how much these methods make use of simple volume overlap over chemical similarity. Additionally, in cases where there are multiple training systems containing the same protein and ligand but with the ligand binding in different conformations, it would be interesting to know if these methods choose one pose to output or if they combine substructures from different poses. By providing all the prediction files and similarity scores against the entire training set, we hope to enable such analyses and drive more granular interpretation of method performance.

We note that the only requirement to apply the Runs N’ Poses benchmark in its entirety on a new method is a structural training cutoff of 30 September 2021, with enough data also available to assess later cutoffs (Figure 2). As the dataset was constructed in an automated fashion, there may be a few complexes which do not qualify as biologically relevant. We expect that careful inspection and continued use will reveal such cases, but since we demonstrate the robustness of the trends discussed across various stratifications, we do not expect this to alter the conclusions.

The future of benchmarking for deep learning methods, as we move toward more complex multidimensional tasks such as cofolding, requires different measures to assess leakage and difficulty, whether for protein-protein, protein-ligand, or nucleic acid-containing complexes. This is also particularly crucial for tasks with even more limited data, such as those involving covalent bonds and modified residues. In this context, we look forward to input from the community on how to best support innovation in this field and strive towards more accurate methods, along with fairer ways of assessing their capabilities.

## 4 Data availability

We provide the entire dataset for benchmarking use at doi.org/10.5281/zenodo.14794785, including the sequences, SMILES, and MSAs used as input for each prediction, and the ground truth CIF and SDF files for each system. For each method, we provide the prediction CIF files, CSV files containing the prediction accuracies (ligand RMSD, LDDT-PLI, pocket backbone RMSD, and LDDT-LP), and confidence scores (ranking score, ligand-protein and protein-ligand chain pair iPTM) for all 25 models across all systems. Further, we provide a CSV file annotating systems with all similarity metrics to the training system with the highest SuCOS-pocket similarity, and the similarity bin that the system falls into, for the September 2021 training cutoff. We also provide a Parquet file with all the similarities found to every system in the PDB up to 2024-06-05, enabling assessment of later training cutoffs. Scripts to recreate the figures in this manuscript, prepare inputs for the different methods, and evaluate predictions can be found at github.com/plinder-org/runs-n-poses. An ML-ready version of the benchmark can be found on Polaris Hub [49] at https://polarishub.io/benchmarks/plinder-org/runs-n-poses. All predicted models along with associated metadata will be made available in ModelArchive [50] with the project identifier ma-rnp.

## 5 Methods

### 5.1 Dataset curation

We ran the PLINDER ingestion pipeline [28] on all entries in the Protein Data Bank (PDB) [51] released after 30 September 2021 until 9 January 2025 to obtain PLINDER systems, defined as all chains from a bioassembly within 6Å of a ligand chain. The resulting systems were further filtered to pass the following criteria:

- Solved by X-ray crystallography
- From the first bioassembly
- With at most 5 ligand chains defined within the system and at most 5 protein chains within the bioassembly
- No systems with covalent ligands
- No systems with all ligands marked as cofactors or oligopeptides/nucleotides/saccharides
- Passing the following X-ray validation criteria (as defined for the PLINDER

benchmark dataset) to ensure that they are reliable ground truth for evaluation:

– Resolution *≤* 3.5Å,

– R-factor *≤* 0.4,

– *R_free_ ≤* 0.45,

– *R − R_free_≤* 0.05,

– Coordinates exist for all heavy atoms in the ligand and pocket,

– No alternative configurations in the ligand and pocket,

– No clash outliers for the ligand,

– No crystal contacts detected for the ligand.

- Passing the following statistics criteria to ensure that they are relevant ligand interactions:

– Having between 3 and 50 detected PLIP [52] interactions,

– Having between 5 and 100 protein residues within 6Å of the ligands in the system,

– The biggest ligand in the system having a molecular weight between 200 and 800.

Systems passing these filters were further clustered to 80% sequence identity, and the system redundancy cluster was defined as the combination of the sequence cluster and the set of CCD [53] codes of the “proper” ligands in the system, i.e ligands not labelled as ions or artifacts (as defined in PLINDER [28]). Only one system per redundancy cluster was chosen, the one with the maximum number of PLIP [52] interactions detected.

These filtering steps resulted in 2,600 systems used as inputs for the four prediction methods (Table 1). For systems where there were polymer chains present in the bioassembly which were not part of the system definition, we used all protein chains in the bioassembly and all ligand chains part of any PLINDER system involving that bioassembly as input for the four methods. While all ligands in the pocket were used for modelling (including ions), the accuracy metrics of only proper ligands were analysed, only for the ligand and protein chains defined for the ground truth PLINDER system, taking the best-scored when multiple mappings were found. Results are presented per proper ligand and no aggregation was done for multi-ligand systems, resulting in 3,047 proper ligands across all the analysed systems. 21 systems contained nucleic acid chains close to the ligand pocket. While cofolding models can handle such input, the chains are currently excluded from PLINDER system definitions and thus would not be scored as part of the system. Future PLINDER releases with fixed system definitions for nucleic acid chains will solve this issue but due to the scarce numbers we chose to drop these 21 systems for the presented analysis. A further 72 system ligands from 63 systems in the common subset could not be analysed as they could not be read with RDKit, and thus similarity scores could not be obtained. For some systems, the SMILES for the ligand in the PDB does not contain chiral information while the ground truth has at least one chiral center, i.e the molecule given as input to each cofolding method may not be the molecule expected as the output. This is not a straightforward issue to fix in an automated fashion, as the ground truth may not have all the atoms resolved and so cannot be the source of truth for extracting the input SMILES. While technically in Runs N’ Poses we ensured that all atoms are resolved, this is not true for many PDB entries, indicating that these methods are also trained on such cases where the chirality information in the input does not match the output. To define prevalent ligands, the number of training systems with ligand analogs was calculated for each system by counting systems before the training cutoff containing ligands with *>*0.9 ligand topological fingerprint Tanimoto similarity [32].

### 5.2 Running all-atom prediction methods

Following the protocol of the PoseBusters benchmark conducted by the authors of AlphaFold3 [22], AlphaFold3 (v3.0.0), Boltz-1 (v0.4.1), Boltz-1x (v1.0.0), Boltz-2 (v2.0.3) Chai-1 (v0.5.1) and Protenix (v0.3.4)were run on one A100 40 GPU each with 10 recycling steps, 200 diffusion sampling steps, 5 seeds and 5 diffusion samples per seed, which resulted in a total of 25 models per system for each method. We use the ligand-protein chain-pair iPTM score for ranking the 25 models, although using the default ranking score showed very similar results.

The input files were prepared according to the instructions for each method. Scripts for input file preparation can be found in our associated GitHub repository. To ensure that the prediction outcomes had no reliance on the depth or quality of the multiple sequence alignments (MSAs) generated by different pipelines, we used the same MSAs as input for all methods. The standard AlphaFold3 MSA generation pipeline was run to obtain the non paired and paired MSAs for each system. These were given as is to Protenix in .a3m format with the pairing and non pairing options, with pairing db set to uniprot as this is the format AlphaFold3 uses for their paired MSA. For Chai-1, both paired and non-paired a3m files were used to generate the aligned.pqt using the provided a3m-to-pqt function, with taxonomy identifiers extracted from UniProt for the paired MSA. The same pairing keys were used to generate the custom MSA CSV files for the Boltz methods. For unpaired sequences, pairing keys were set to −1. Thus, each method uses the same MSAs and pairing strategies as input. In addition, Chai-1 predictions were made with the ESM embedding option enabled and AlphaFold3 predictions were made with templates enabled. AlphaFold3 was also run without templates for comparison (AF3-NT). For RosettaFold-All-Atom, we followed the standard procedure as described in its documentation.

We did not manage to obtain prediction results from all methods across all 2,600 systems in the dataset. Some systems (128) failed to produce output across all four methods due to (1) incorrect specification of modified amino acids, (2) reaching memory limits in A100 40G GPUs, and (3) failures in the conformer generation step. Table 1 details the number of systems which could be modelled by all the different methods, with Boltz-1/Boltz-1x having the highest number of missing predictions. In total we had 229,887 predictions across the four methods used for common subset analysis, namely AlphaFold3 (with templates), Boltz-1, Chai-1, and Protenix, resulting in a common subset of 2,311 proper ligands (3,288 including ions) across 2,077 systems. Note: Boltz-2 was only run on 1,028 systems released after its training cutoff of 2023-06-01.

### 5.3 Accuracy scoring

To evaluate the accuracy of predicted models, we use the compare-ligand-structures action of OpenStructure [54] (version 2.8.0), which returns the binding-site superposed symmetry-corrected ligand RMSD (hereafter referred to just as RMSD), the local difference distance test of the ligand pocket (LDDT-LP), and the local difference distance test of protein-ligand interactions (LDDT-PLI). For LDDT-PLI calculation we enable the “added model contacts” flag which further penalises protein-ligand contacts present in the model which are not in the reference (we note that this is the default for LDDT-PLI since OpenStructure 2.9). In addition, we calculated the F1-score of pocket recovery, where true positives are defined as pocket residues found in both the predicted and ground truth structures (where pocket residues are defined as those having any atom within 6Å of any ligand atom). All of these scores are calculated per-ligand using the receptor CIF and ligand SDF files saved by PLINDER as the ground truth for each system. Only the scores for ligands labelled as proper ligands (i.e excluding ions and artifacts) were analysed, and multi-ligand systems form multiple data points in our analysis, one per proper ligand. We provide code to extract relevant scores from the OpenStructure output JSON files in our associated GitHub repository.

As we are working with cofolding methods, where both the protein pocket and ligand pose have to be accurately modelled, we redefine the commonly used “Success rate” measure as the fraction of systems with *<*2Å RMSD and *>*0.8 LDDT-PLI. Figure 7A shows the distribution of RMSDs and LDDT-PLIs across all predictions. This addresses two kinds of outliers on the top right and the bottom left.

**Fig. 7.**
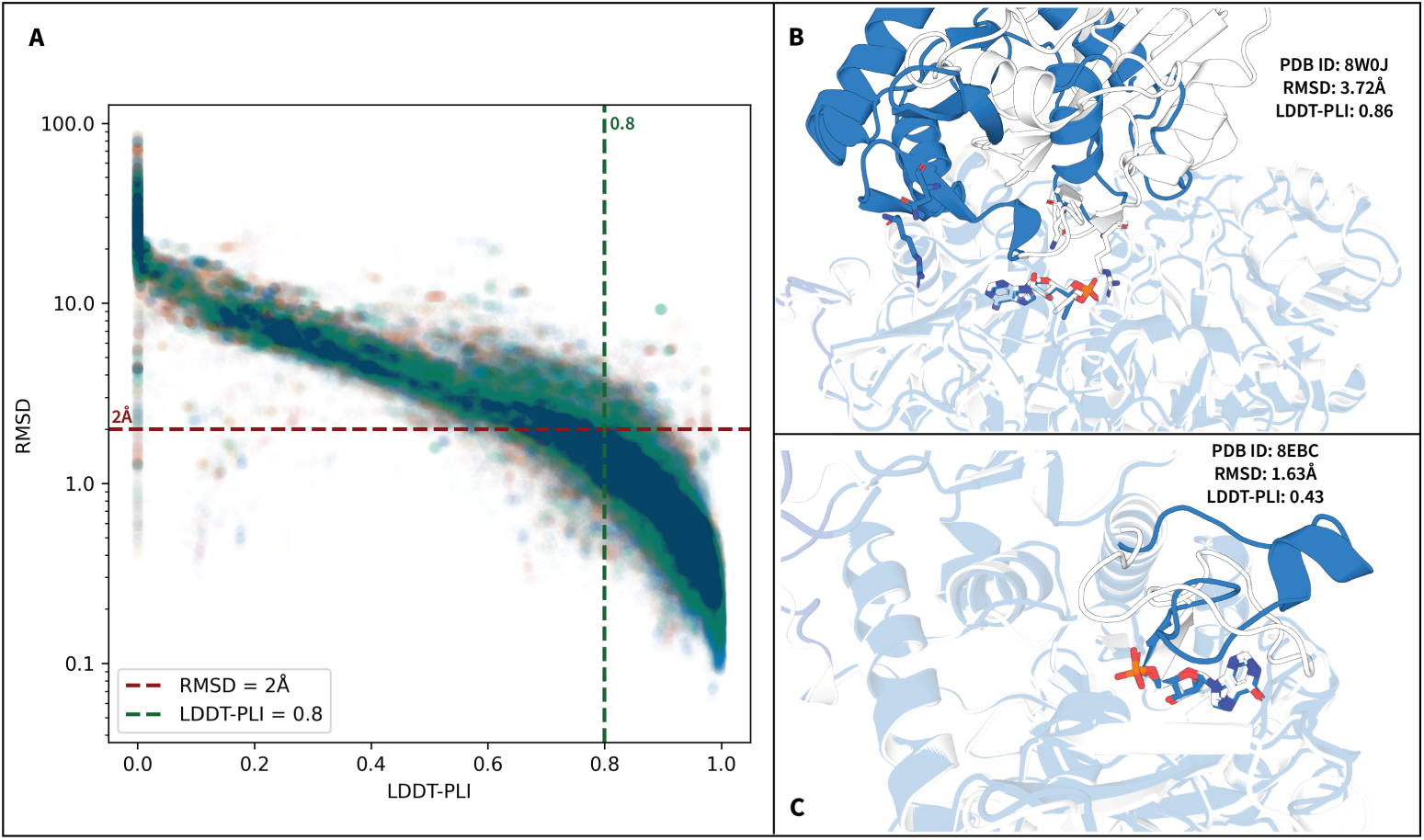
LDDT-PLI and RMSD scores across all predictions. **A)** The distribution of binding site superposed RMSD (log-scale) and LDDT-PLI for all 229,887 predictions with 2Å RMSD and 0.8 LDDT-PLI shown as dashed lines. **B)** An example with high RMSD and high LDDT-PLI, the binding site residues used for performing the superposition are shown as sticks in the predicted structure (blue) and ground truth (gray). **C)** An example with low RMSD and low LDDT-PLI, prediction in blue and ground truth in gray.

The top right has 1.6% of predictions with LDDT-PLI *>* 0.8 but RMSD *>* 2Å, sometimes consisting of cases such as the example shown in Figure 7B. While the global superposition here would have resulted in a low RMSD, and the LDDT-PLI is high (0.86) due to many of the protein-ligand distances with one domain being maintained, the binding site superposed RMSD is high 3.72Å as the two binding site residues shown in sticks are used to perform the superposition and the top domain is incorrectly modelled. This area also has some cases where a part of the ligand not interacting with the protein is flexible and therefore its orientation is not biologically relevant nor well-resolved. While RMSD might correct the overoptimistic results of LDDT-PLI in the former case, in the latter case LDDT-PLI might be a more meaningful measure on its own. However, as *<*2% of cases across all 229,887 predicted models fall into this area and we cannot automatically differentiate between the two cases, we opted to treat these as failures.

The bottom left has 8.3% of the predictions with RMSD *<* 2Å but LDDT-PLI *<* 0.8, consisting of cases where the pose is correct but the pocket is wrongly modelled.

The example in Figure 7C shows the global superposition of the two complexes, which would lead to a very low RMSD value. While the binding-site superposed RMSD shows worse results (1.63 Å), it would still be considered as a success if only using RMSD *<*2Å as the success criteria. LDDT-PLI on the other hand is 0.43, correctly reflecting that the protein-ligand interactions are not maintained.

To measure physical plausibility of the predicted poses, we ran the suite of checks implemented in PoseBusters [19] using the “dock” configuration on all predicted models, and report the number of systems that pass all default checks.

### 5.4 Similarity scoring

PLINDER calculates 14 metrics covering protein sequence (similarity and identity), protein structure (CA-LDDT); pocket coverage, identity and structure; and the similarity of PLIP interactions [28]. In addition, the Combined Overlap Score (SuCOS), which measures the overlap of volume and chemical features between two ligand poses, has been shown to better differentiate binding pose similarity compared to RMSD and PLI-based scores [30]. Thus, we calculated three more metrics working in concert with the protein-based scores.

As SuCOS calculation requires a superposition between ligand poses, we align the ligand poses of a query and target system using RDKit’s rdShapeAlign.AlignMol [55] command. The superposed molecules were then used to calculate the SuCOS score [29], as implemented in https://github.com/susanhleung/SuCOS. These three scores are further multiplied by the pocket coverage (pocket qcov, divided by 100 to get the fraction), which measures the percentage of residues within 6Å of the ligand in the query system which align with residues within 6Å of the ligand in the target system, to obtain relevant scores measuring the similarity of ligand poses within the same pocket. While approaches combining protein and pocket-level superposition were attempted, they turned out to be too sensitive to the efficacy of rigid protein superposition.

The main similarity score used in this manuscript to define training set similarity, find the closest training system as well as cluster similar systems, unless otherwise specified, is the SuCOS multiplied by the pocket qcov (SuCOS-pocket). To obtain these values for each system in our set, we perform a Foldseek [56] search against the entire PDB and calculate similarities between any pair of test and train systems where any protein chain has an alignment. Note that pocket qcov does not directly consider residue identity or similarity, and just checks that the residues are aligned.

Despite our best efforts, we cannot be sure that we are always able to accurately calculate similarity to the training set, as our methods rely on a combination of alignment and superposition which may fail in edge cases. Also, we only search for similarity against PLI systems, excluding only-artifact systems and those with oligopeptides, oligosaccharides or oligonucleotides over 10 in length, thus potentially missing cases of training set proteins complexed with oligo, protein or nucleotide chains which resemble small molecule binding. However, this latter similarity would indicate transfer of binding modes from different modalities and perhaps may not be considered memorisation.

Using PLINDER’s pli qcov scores, which counts the fraction of shared protein-ligand interaction types detected by PLIP [52] on aligning residues, gave similar results (Supplementary Figure S3L) - however, as cofolding methods are not trained to recapitulate interactions, we wanted to account for cases where the SuCOS similarity of the ligand pose remains high but pocket residue conformational changes result in different PLIs being detected (i.e low pli qcov similarity, high SuCOS-pocket similarity), as well as cases where the part of the ligand interacting with the protein remains the same, but other regions of the ligand change in conformation (i.e high pli qcov similarity, low SuCOS-pocket similarity).

The dataset was clustered using community clustering on the graph connecting systems having SuCOS-pocket similarity scores above 50. Cluster representatives are chosen as those with the lowest similarity to the training set.

Supplementary Figure S4 shows examples of the kinds of ligands found in each training set similarity bin.

## Acknowledgements

We thank the members of the Schwede group for technical support, useful suggestions, and text revisions, Vladas Oleinikovas, Lukas Jarosch, Seohyun (Chris) Kim for insightful discussions and comments, Cas Wognum for the Polaris set-up, GitHub commenters for spotting and reporting minor errors in the previous versions of the manuscript, and sciCORE at the University of Basel (https://scicore.unibas.ch/) for providing computational resources and storage space. We gratefully acknowledge financial support for parts of this work by the SIB Swiss Institute of Bioinformatics (https://www.sib.swiss/), the Biozentrum of the University of Basel (https://www.biozentrum.unibas.ch/), the Swiss National Science Foundation (SNSF; Ambizione grant 223634), and Dompé farmaceutici. This work uses outputs from AlphaFold3, subject to the AlphaFold 3 Output Terms of Use (version 3.0.0 [commit 2ffe43f], accessed on 15 Nov. 2024). All AlphaFold3 calculations and analyses were carried out by and using the weights provided to Jérôme Eberhardt.

---

**Supplementary Figure S1.**
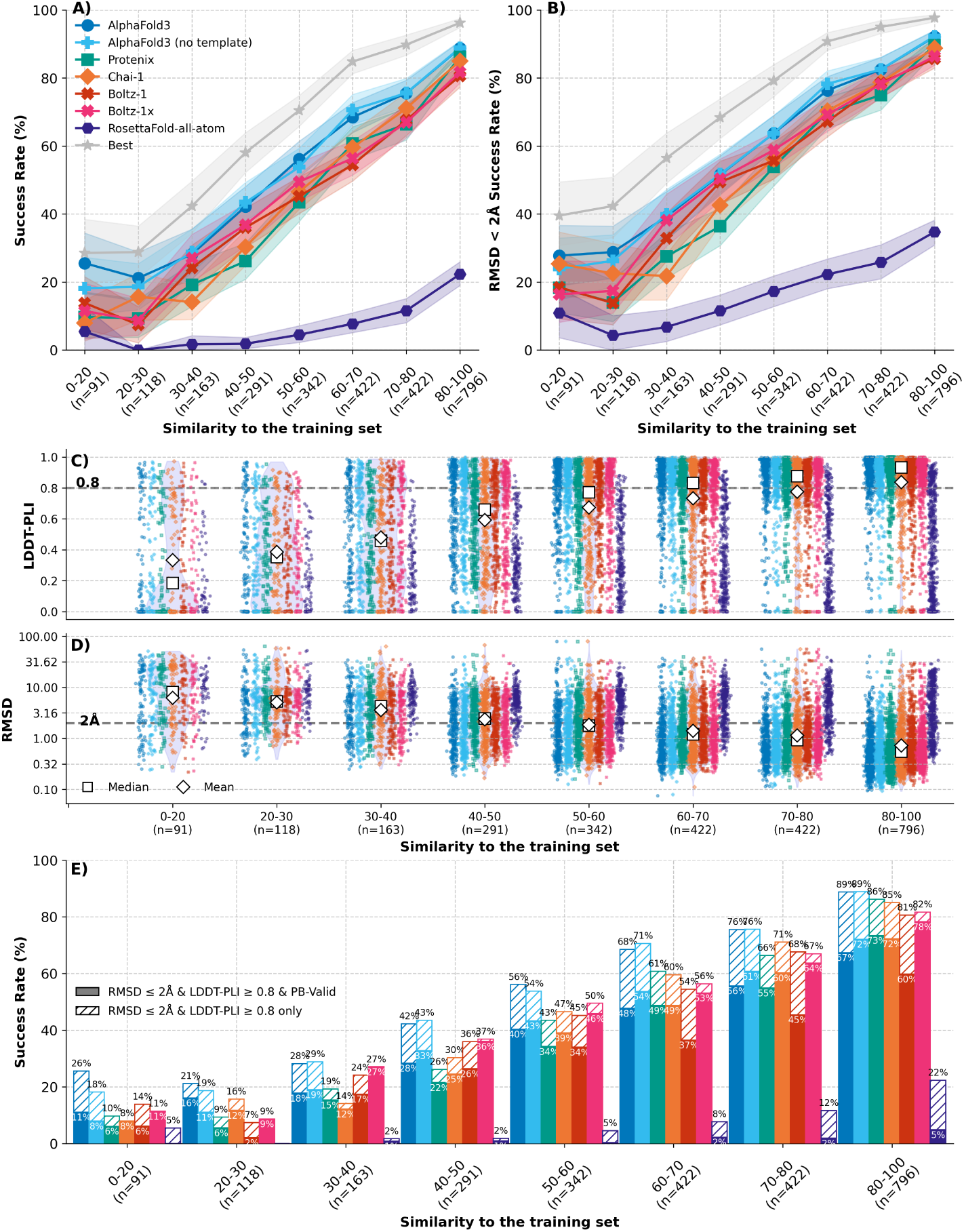
Prediction accuracy vs. training set similarity across all predicted models. **A)** The success rate, defined as the percentage of system ligands with *<*2Å RMSD and *>*0.8 LDDT-PLI, for all system ligands in each similarity bin across the assessed methods and when selecting the best-scored model from the top-ranked. Shaded regions correspond to the 95% confidence interval, calculated from 1,000 bootstrap samples for each bin and method. **B)** same as A but with success rate defined only by *<*2Å RMSD. **C), D)** The distribution of LDDT-PLI and RMSD values respectively for all methods displayed as a violin plot, with each individual method being shown as coloured scatter points. **E)** Same as panel A with added physical validity checks. The striped bars show the share of predictions of each method that have *<*2Å RMSD and *>*0.8 LDDT-PLI and the solid bars show the subset that in addition have valid geometries and energies, i.e., pass all PoseBusters tests and are therefore ‘PB-Valid’.

**Supplementary Figure S2.**
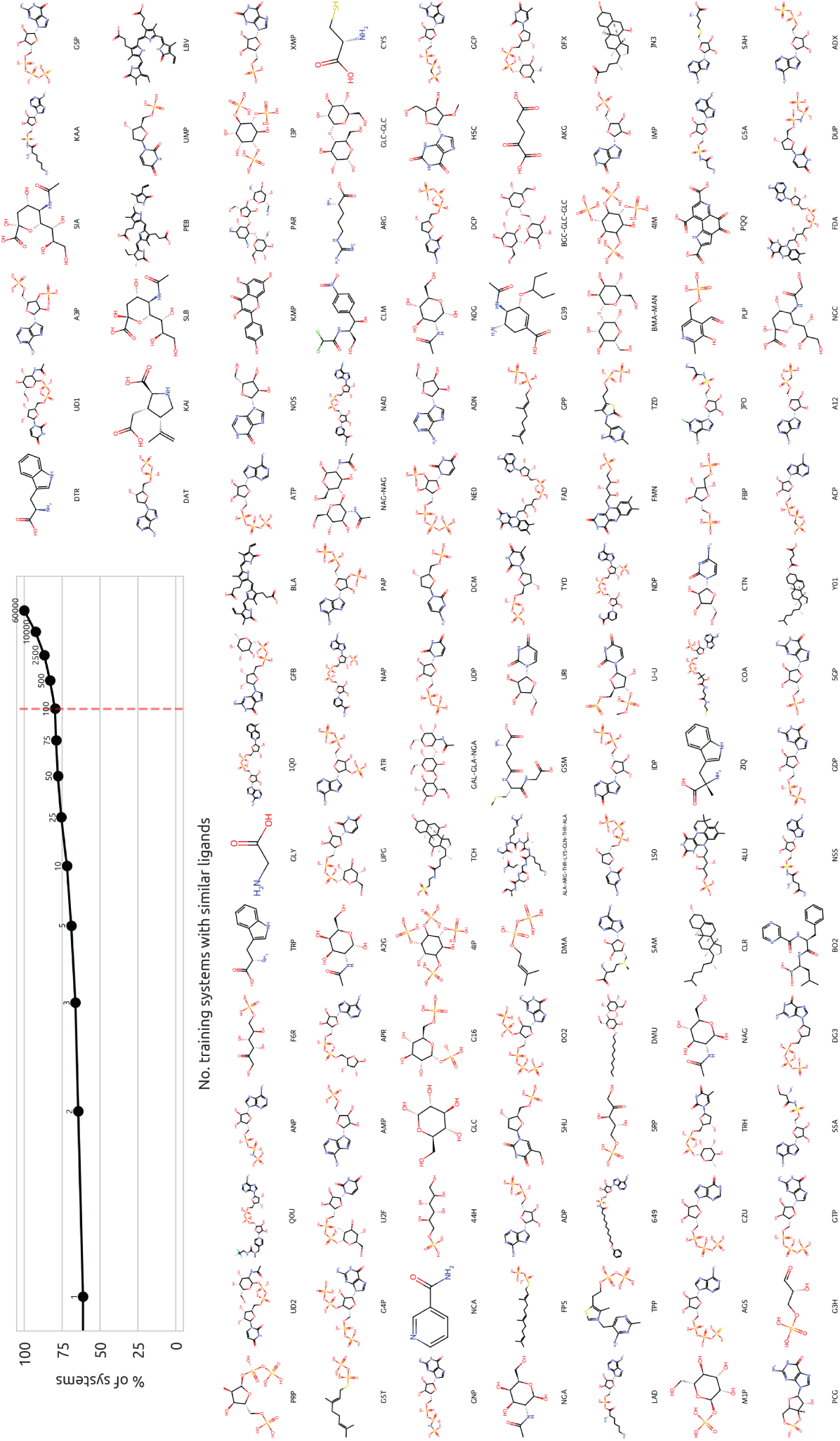
Prevalent ligands. The percentage of system ligands (Y-axis) having different numbers of training systems with analogous ligands (X-axis, log scale), and all the ligands within systems with *>*100 analogous ligand training systems.

**Supplementary Figure S3.**
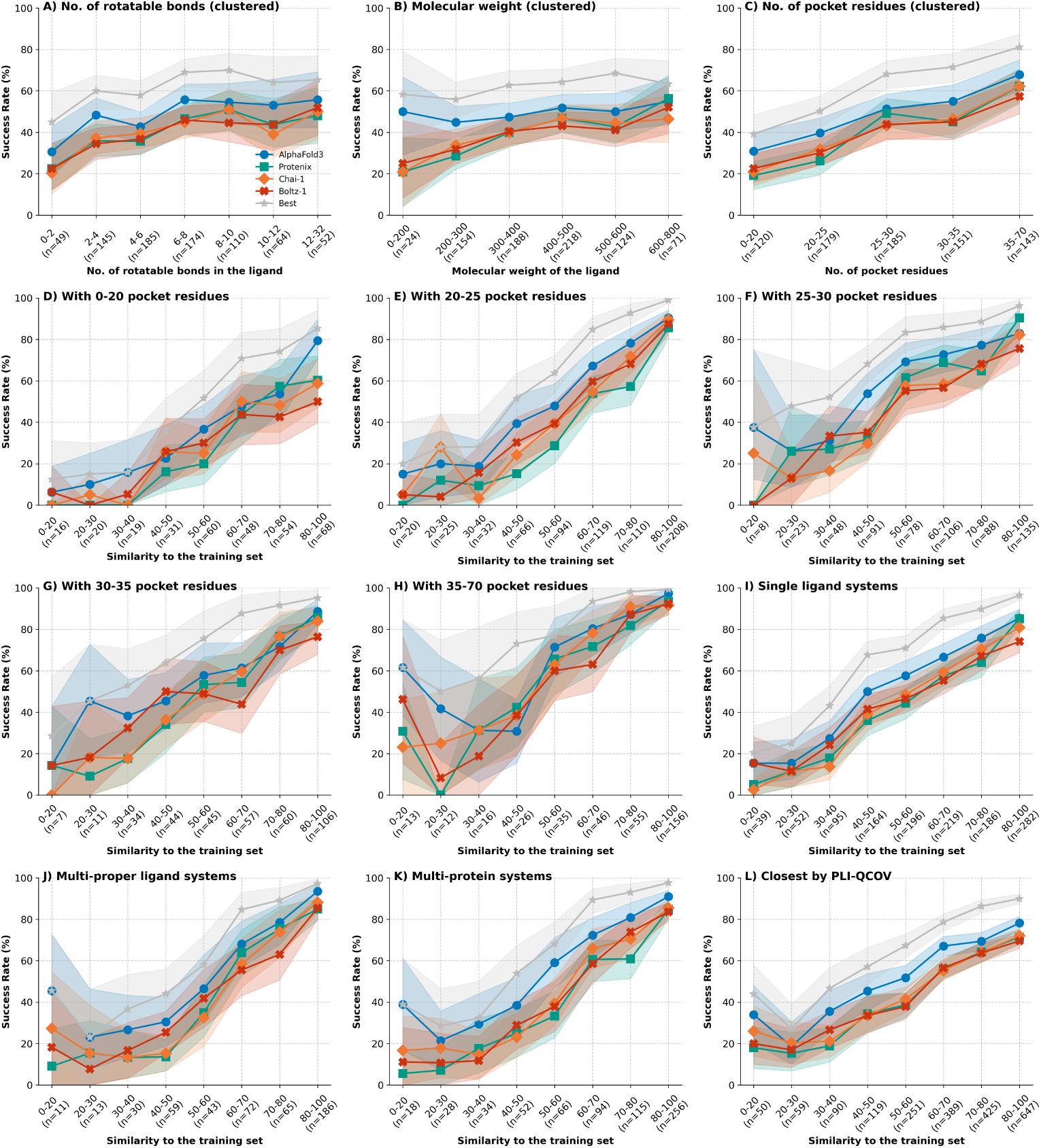
Success rate distributions for different stratifications. Success rates for the different methods and the best model of the four, **A)** for clustered system ligands with differing numbers of rotatable bonds. **B)** for clustered system ligands with differing molecular weights. **C)** for clustered system ligands with differing numbers of residues in the binding pocket. **D-H)** or system ligands having binding pockets (residues within 6 Å of the ligand) defined by D) 0-20, E) 20-25, F) 25-30, G) 30-35, and H) *>*35 residues respectively. **I-K)** across the different SuCOS-pocket similarity bins only for systems having I) a single ligand, J) multiple proper ligands, and K) multiple protein chains, respectively. **L)** across different pli qcov similarity bins when using pli qcov similarity to find the closest training system.

**Supplementary Figure S4.**
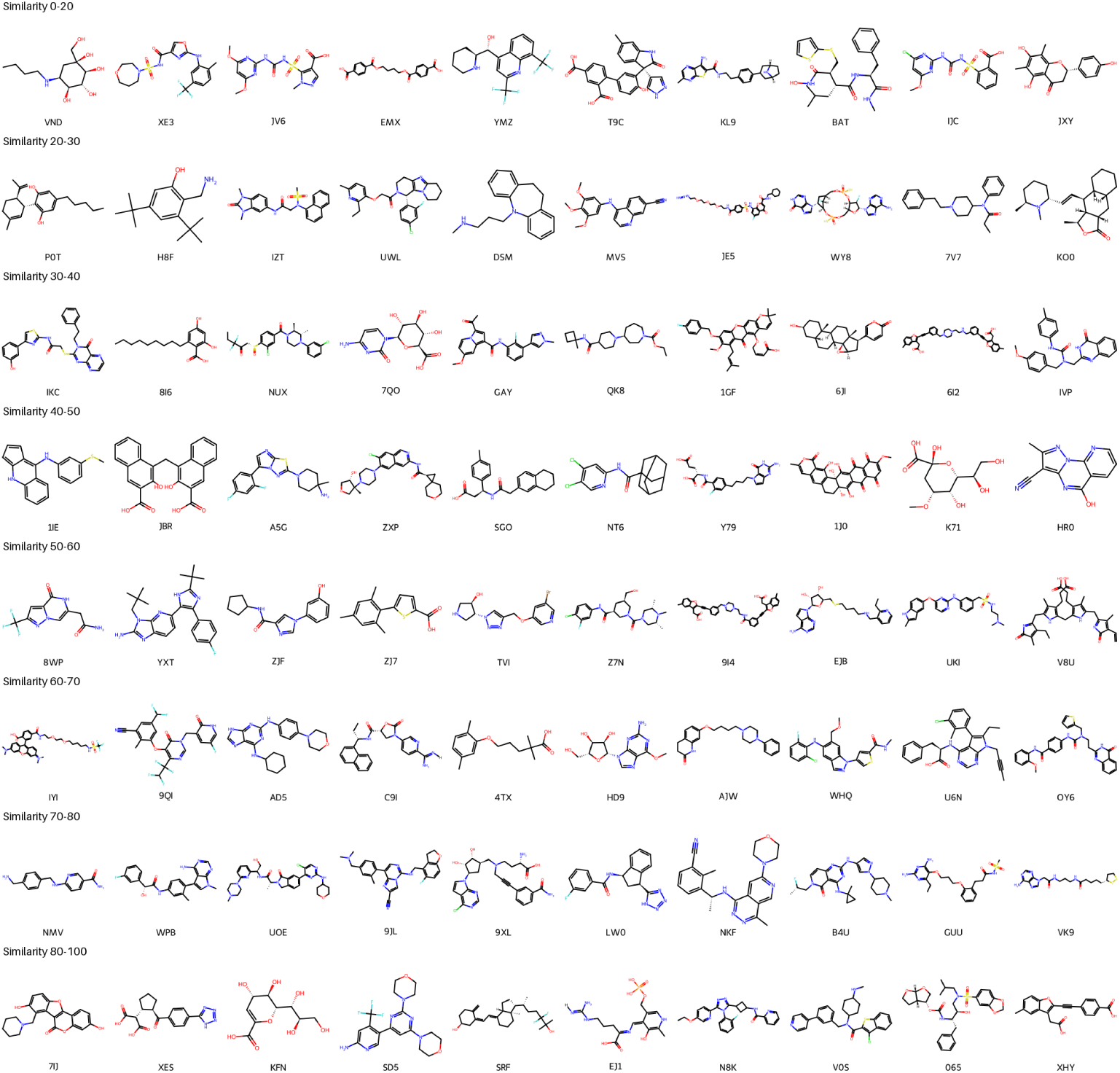
Ten examples of small molecules found in each training set similarity bin.

**Supplementary Table S1.** Representatives of populated clusters. Sizes of the ten most populated clusters, along with the PDB ID and description of the cluster representative with the lowest training set similarity for each, and distribution of the cluster across different training SuCOS-pocket similarity bins.

| Index | Cluster Size | PDB ID | Description | Similarity bins |
| --- | --- | --- | --- | --- |
| 1 | 171 | 7EN9 | Crystal structure of SARS-CoV-2 3CLpro in complex with the non-covalent inhibitor WU-02 | 80-100 (79), 70-80 (46), 60-70 (25), 50-60 (15), 40-50 (6) |
| 2 | 136 | 8OKU | Salt-Inducible Kinase 3 in complex with an inhibitor | 60-70 (38), 70-80 (37), 80-100 (28), 50-60 (19), 40-50 (11), 30-40 (3) |
| 3 | 100 | 5SH0 | Crystal structure of human phosphodiesterase 10 in complex with IZT | 50-60 (33), 40-50 (26), 30-40 (16), 60-70 (13), 80-100 (5), 20-30 (4), 70-80 (3) |
| 4 | 92 | 8HUU | Crystal structure of HCoV-NL63 main protease with S217622 | 80-100 (43), 60-70 (20), 70-80 (20), 40-50 (5), 50-60 (4) |
| 5 | 73 | 7S3Z | Structure of rat neuronal nitric oxide synthase heme domain in complex with VOP | 80-100 (51), 70-80 (16), 60-70 (5), 50-60 (1) |
| 6 | 43 | 8JOO | Crystal structure of cytochrome P450 IkaD from *Streptomyces* sp. ZJ306, in complex with the substrate ikarugamycin | 80-100 (23), 60-70 (5), 70-80 (5), 30-40 (4), 40-50 (3), 50-60 (3) |
| 7 | 35 | 7G00 | Crystal structure of human FABP1 in complex with WII | 60-70 (13), 50-60 (10), 40-50 (6), 70-80 (3), 80-100 (2), 30-40 (1) |
| 8 | 28 | 7UBO | Crystal structure of the first bromodomain of human BRDT in complex with the inhibitor CCD-956 | 70-80 (7), 80-100 (7), 50-60 (6), 60-70 (5), 40-50 (2), 30-40 (1) |
| 9 | 28 | 7S1S | PRMT5/MEP50 crystal structure with MTA and MRTX-1719 bound | 80-100 (12), 40-50 (6), 50-60 (3), 60-70 (3), 30-40 (2), 70-80 (2) |
| 10 | 26 | 7V0H | Crystal structure of putative glucose 1-dehydrogenase from *Burkholderia cenocepacia* in complex with NADP and a potential reaction product | 80-100 (13), 60-70 (7), 70-80 (6) |

## Notes

https://github.com/plinder-org/runs-n-poses

https://doi.org/10.5281/zenodo.14794785

### Competing Interest Statement

The authors have declared no competing interest.

### Summary of Updates

- Include Boltz-2 and benchmarking newer cutoffs - Posebusters physical plausibility checks - Include AF3-no template, RosettaFold-all-atom, Boltz-1x results - Added analysis of classification accuracy of iPTM-based confidence metrics - Added binding pocket detection accuracy - Expanded text

